# SpatialData: an open and universal data framework for spatial omics

**DOI:** 10.1101/2023.05.05.539647

**Authors:** Luca Marconato, Giovanni Palla, Kevin A. Yamauchi, Isaac Virshup, Elyas Heidari, Tim Treis, Marcella Toth, Rahul B. Shrestha, Harald Vöhringer, Wolfgang Huber, Moritz Gerstung, Josh Moore, Fabian J. Theis, Oliver Stegle

**Author notes:** contributed equally (Luca Marconato, Giovanni Palla, Kevin A. Yamauchi, Isaac Virshup).

## Abstract

Spatially resolved omics technologies are transforming our understanding of biological tissues. However, handling uni- and multi-modal spatial omics datasets remains a challenge owing to large volumes of data, heterogeneous data types and the lack of unified spatially-aware data structures. Here, we introduce SpatialData, a framework that establishes a unified and extensible multi-platform file-format, lazy representation of larger-than-memory data, transformations, and alignment to common coordinate systems. SpatialData facilitates spatial annotations and cross-modal aggregation and analysis, the utility of which is illustrated via multiple vignettes, including integrative analysis on a multi-modal Xenium and Visium breast cancer study.

## Introduction

The function of biological tissues is strongly linked to their composition and organization. Advances in imaging and spatial molecular profiling technologies enable addressing these questions by interrogating tissue architectures with ever growing comprehensiveness, resolution and sensitivity^1, 2^. Existing spatial molecular profiling methods quantify DNA, RNA, protein, and/or metabolite abundances in situ. Several of these technologies employ light microscopy, providing spatial resolution of morphological features at length scales from subcellular to entire organism. Critically, spatial omics technologies are rapidly evolving, and each data modality features distinct advantages and limitations (e.g., spatial resolution, molecular multiplexing, detection sensitivity). The ability to efficiently integrate and then operate on data from different spatial omics modalities promises to be instrumental for constructing holistic views of biological systems.

While progress has been made in analyzing individual spatial omics datasets, integrating multimodal spatial omics data remains a practical challenge (Supplementary Note 1, Table S1). Firstly, loading datasets into analysis pipelines in a coherent manner is hampered by the diversity in data types (e.g., tabular data for sequencing and 10s-100s GB dense arrays for images) and file formats (e.g., technology-specific vendor formats). Additionally, individual spatial omics modalities often have different spatial resolutions and the data can be acquired from different regions of the same tissue. Thus, in order to integrate such data, they must be appropriately transformed to align them to a common coordinate system (CCS). Aligning datasets to a CCS is a building block to establish a global common coordinate framework (CCF)^3^. Finally, untangling the complexity of multimodal spatial omics datasets requires expert domain knowledge, motivating approaches that enable large-scale interactive data exploration and annotation. Thus, to unlock the full potential of emerging spatial multiomics studies^2, 4^, there is a need for computational infrastructures to store, explore, analyze, and annotate diverse spatial omics data with a unified programmatic interface.

## Results

### SpatialData concept and implementation

The SpatialData framework enables the FAIR^5^ integration of multimodal spatial omics data. A language-independent storage format increases the interoperability of data sources, post-alignment, while the Python library standardizes the access of and operation across diverse data types. The SpatialData format supports all major spatial omics technologies and derived quantities (Figure 1A, Figure 1C, Supplementary Note 2, Table S3). Briefly, spatial datasets are represented using five primitive elements: Images (raster images), Labels (e.g. raster segmentation masks), Points (e.g. molecular probes), Shapes (e.g., polygon regions of interests, array capture locations etc.), and Tables (e.g., molecular quantifications and annotations) (Table S2). The file format also keeps track of any coordinate transformation or alignment steps applied to individual datasets. Collections of datasets from multiple assays can be stored within a single SpatialData file and because the spatial relationship between them is stored via coordinate transformations, they can be thus analyzed together. The SpatialData format is built upon the OME-NGFF specifications and leverages the Zarr file format (Supplementary Figure 1), which provides performant, interoperable access for both traditional filesystem-based as well as cloud-based storage^6, 7^.

**Figure 1.**
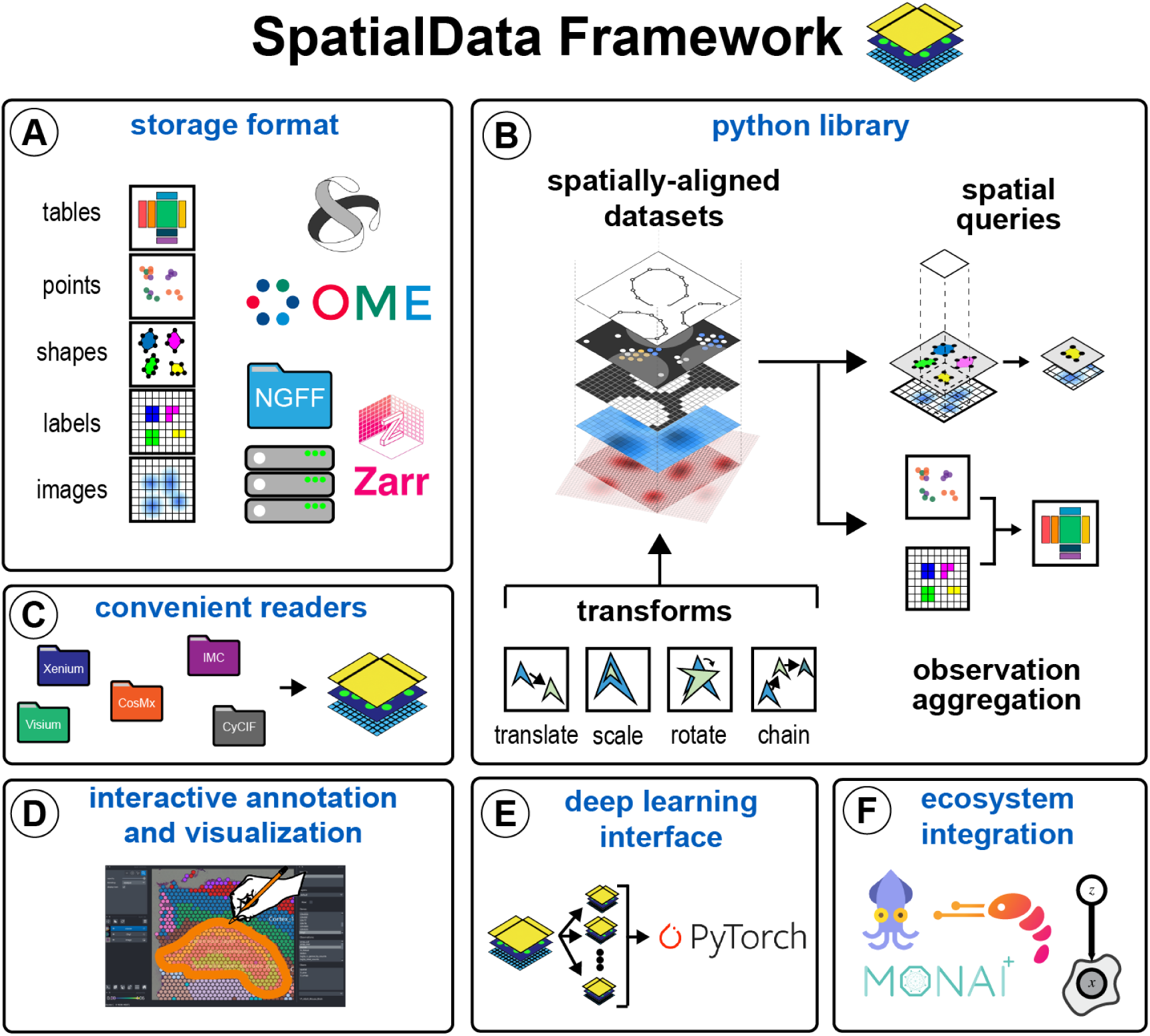
Design overview and core functionality of SpatialData. **(A)** The SpatialData storage format provides uniform storage for raw and derived data of diverse spatial omics technologies. The format builds on five primitive elements (SpatialElements), stored to Zarr in an OME-NGFF compliant manner. **(B)** The SpatialData Python library implements core operations for data access, alignment, queries, and aggregation of spatial omics data. Transformations align multiple SpatialElements to a common coordinate system (CCS). The CCS allows unified spatial queries and aggregation operators to be deployed across datasets. **(C)** SpatialData provides access to various data formats, including vendor-specific file formats. Multiple datasets can be stored in a single file and together are represented as a SpatialData object. **(D)** Datasets stored in the SpatialData format can be annotated interactively using the integrated napari-spatialdata plugin. SpatialData also provides functionality for generating static plots. **(E)** Leveraging an implementation of the PyTorch Dataset class, deep learning models can be trained directly from SpatialData objects. **(F)** Since SpatialData builds upon established standards and software, it ties into the existing ecosystem for multimodal analysis including Squidpy ^8^, Scanpy^9^, MONAI ^10^ and scvi-tools^11^, amongst others.

The SpatialData Python library represents this format as SpatialData objects in memory, which supports lazy loading of larger-than-memory data (Figure 1B). The SpatialData Python library also provides reader functions for current spatial omics technologies (Figure 1C, Table S2). The library provides versatile functionality for manipulating and accessing SpatialData objects. A core feature is the efficient definition of common coordinate systems (CCSs) of biological systems^3^. Briefly, datasets from individual data modalities are associated with coordinate transformations (Figure 1B). SpatialData implements affine coordinate transformations, which can be composed (<u>link to tutorial</u>). Once aligned, datasets can be queried (Supplementary Note 3, Supplementary Figure 2, <u>link to tutorial</u>) and aggregated (Supplementary Note 4, Supplementary Figure 3, <u>link to tutorial</u>) using spatial annotations both within and across modalities. The query and aggregation interface also allows for creating new datasets grouped by biologically-informed factors from large aligned data, thereby facilitating exploration and data access.

SpatialData comes with a napari plugin for interactive annotation (napari-spatialdata) (Figure 1D, Supplementary Figure 4, Supplementary Note 5). The napari-spatialdata plugin allows analysts to make spatial annotations such as drawing regions of interest (<u>link to tutorial</u>) or landmarks for guiding multi-dataset registration (<u>link to tutorial</u>). Static figures and graphics can be created using the spatialdata-plot library (Supplementary Figure 5, Supplementary Note 6. <u>Link to tutorial</u>).

The SpatialData library seamlessly integrates with the existing Python ecosystem by building upon standard scientific Python data types. We have implemented a PyTorch Dataset class so that deep learning models can be easily trained directly from SpatialData objects (Figure 1E, Supplementary Note 7 Supplementary Figure 6). Further, the analysis packages in the scverse ecosystem^12^ can be used to analyze SpatialData objects (Figure 1F, Supplementary Note 8). Taken together, the SpatialData framework provides infrastructure for integrating and analyzing spatial omics data.

### Application of SpatialData to a multi-modal and multi-technology breast cancer experiment

To illustrate SpatialData’s applicability to multi-modal integration and analysis, we applied the framework to a breast cancer study that combines H&E images, 10x Genomics Visium and Xenium assays^13^. The study comprises two *in situ* sequencing (Xenium) and one Spatial Transcriptomics dataset (10x Visium CytAssist) from consecutive sections of a breast cancer tumor. First, we used napari-spatialdata to define landmark points that are present in all datasets. We then aligned all three datasets using affine transformations, thereby defining a CCS for this dataset (Figure 2A). As a result of the alignment, SpatialData enabled us to identify the common spatial area, which can be accessed using SpatialData queries across all three datasets.

**Figure 2.**
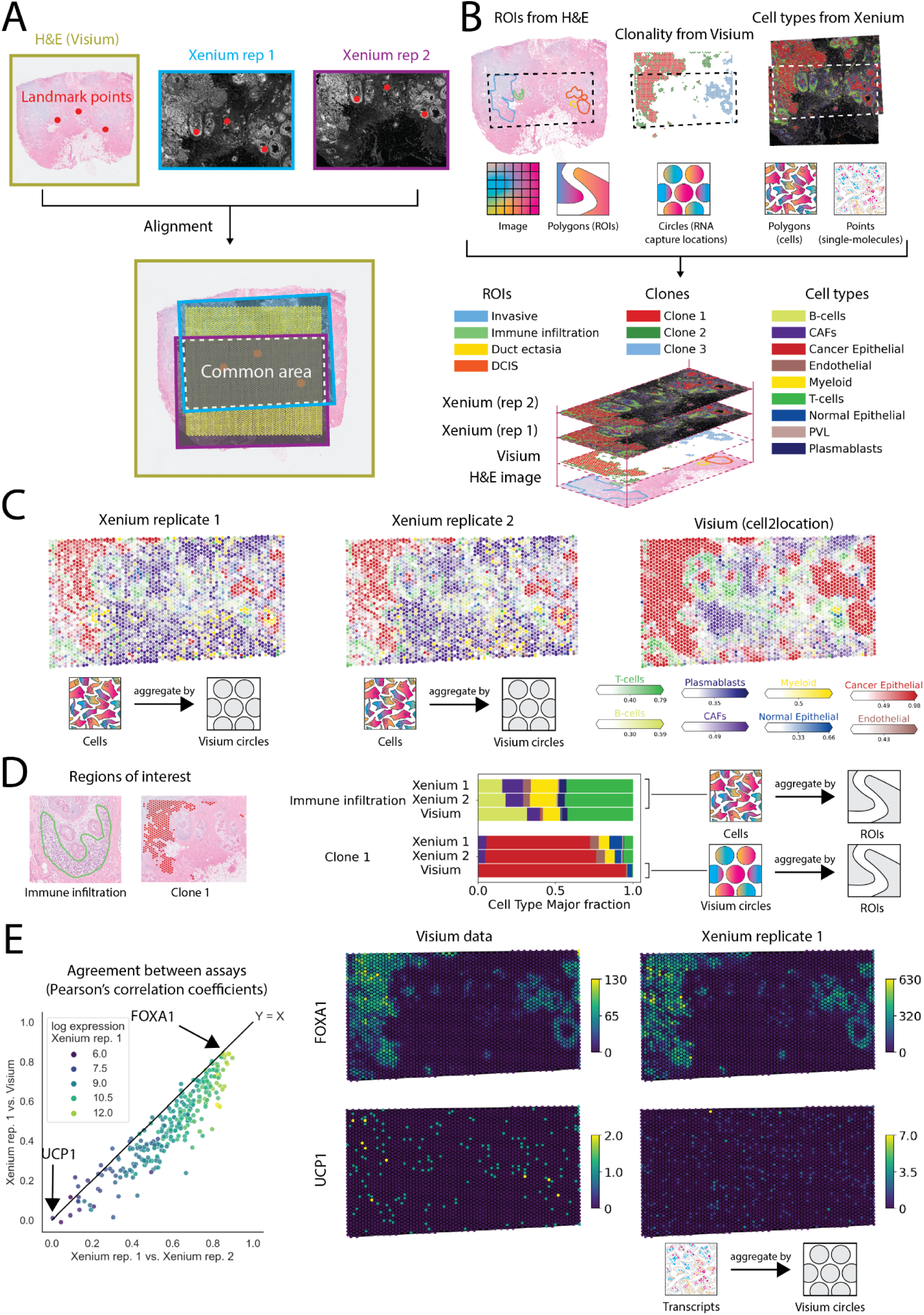
Alignment and integrative analysis of three spatial datasets. **(A)** Registration of two breast cancer Xenium replicate slides, one Visium slide, and their corresponding H&E images to a common coordinate system (CCS) based on interactively selected landmarks. **(B)** Spatial annotations can be transferred across datasets using the CCS. Top: spatial annotations derived from multiple datasets, including histological regions of interest (H&E image), tumor clones (Visium-derived copy number aberrations) and cell types (Xenium and scRNA-seq). Spatial annotations are represented by different spatial elements (polygons, circles, molecules) and can be transferred between datasets via the CCS. The visualizations are presented in a larger format in Supplementary Note 11, Supplementary Figure 10. **(C)** SpatialData queries facilitate cross-modality aggregation for data integration, quality control and benchmarking. Left, middle: Cell-type fractions in Xenium computed at circular regions that correspond to Visium quantification locations. Right: Cell-type fraction estimates from deconvolution methods based on Visium data (using cell2location). **(D)** SpatialData queries allow for arbitrary geometrical quantifications. Shown are cell-type fraction estimates obtained in Xenium or Visium (via spot-level cell2location estimates) at annotated ROIs and clones as in **B. (E)** Comparison of gene expression quantification in Xenium and Visium using SpatialData aggregations at Visium capture locations. Left: Scatter plot of the correlation coefficient of aggregated gene expression quantifications between Xenium replicates (x-axis) versus the correlation coefficient between Xenium and Visium (y-axis). Shown are gene-expression quantifications for 313 genes present in both Xenium and Visium. Point color denotes log expression in Xenium replicate 1. Lowly expressed genes are associated with reduced correlation. Right: Visualization of aggregated expression levels at Visium locations for FOXA1 (top) and UCP1 (bottom). Color bars denote raw counts.

Next, we used the collective information from all three datasets to create a shared set of spatial annotations. We selected four regions of interest (ROIs) based on histological features present in the H&E image using napari-spatialdata. We then used genome-wide transcriptome information in Visium to infer copy number variations (using CopyKat ^14^) and select major genetic subclones. Finally, we annotated the Xenium replicates using a label transfer method (ingest, implemented in scanpy^9^), using cell type labels of an independent breast cancer scRNA-seq atlas^15^ (Figure 2B, Supplementary Figure 8B).

To exemplify how SpatialData can be used to transfer spatial annotations, we considered the Visium capture locations and quantified cell type fractions in each location by aggregating the cell type information from the overlapping Xenium cells. For comparison, we also considered a deconvolution-based analysis of Visium using cell2location^16^ in conjunction with the same scRNA-seq-derived cell types^15^ as reference. We observed high concordance between Xenium replicates (median Pearson’s R=0.75 across Visium locations) and overall good agreement between Xenium and deconvolution-based estimates (median Pearson’s R=0.60).

Analogously to the aggregation at Visum locations, we considered ROIs defined from H&E and the areas defined by the union of the subclone locations from Visium (Figure 2D, Supplementary Figure 8A). Again, we quantified cell types within each region, either directly using cell count fractions from Xenium or via deconvolution of the corresponding Visium data. Again, the two Xenium replicates showed high concordance of cell type fractions, and the Xenium and Visium were consistent. Collectively, these examples illustrate how the SpatialData aggregation functions can be used to integrate signals between samples, and how the framework facilitates quality assessment between replicates, and benchmarking of computational methods.

As a second aggregation use case, we compared expression quantifications of individual genes measured by the Xenium and Visium assays. To do so, we aggregated individual molecule counts from the Xenium data at Visium capture locations. We compared transcript-to-circles aggregations from the Xenium replicates to the original Visium transcript counts (Figure 2E, Supplementary Figure 8B). As expected the aggregated counts were highly concordant between the Xenium replicates (median Pearson’s R=0.62, Figure 2E, Supplementary Figure 8C-E) and to a lesser extent between a Xenium and Visium counts (median Pearson’s R=0.48, Supplementary Figure 8C-E). We also observed a direct relationship between the overall transcript abundance and the agreement between different tissue sections and technologies (Figure 2E).

In sum, these examples illustrate the flexibility of the aggregation functionality, which can be applied between SpatialElements of different kinds (points, circular capture locations, cells, larger anatomical ROIs) to transfer diverse types of spatial annotations (cell expression, cell types fractions). Further examples and advanced use cases of SpatialData aggregation operations are discussed in Supplementary Note 4.

### Application to further datasets and illustration of additional use cases

SpatialData is useful for processing single modality datasets in addition to complex multi-modality studies as shown above. The SpatialData framework comes with additional vignettes that illustrate other use cases. We illustrate how SpatialData can serve as a backend to facilitate the training of deep learning models (Supplemental Note 7, <u>link to</u> <u>tutorial</u>) and popular spatial interpretation tools such as Squidpy (l<u>ink to tutorial</u>). As a starting point for using SpatialData in conjunction with different technologies, we also provide pre-formatted SpatialData objects from over 40 datasets across a total of 8 technologies (Table S3). Interactive annotation can be performed on both single- and multi-modality datasets (l<u>ink to tutorial</u>, <u>link to tutorial</u>). Finally, we demonstrate how SpatialData can be used to align multiple fields of view to a global reference coordinate system, by mapping 12 Visium slides to a large prostate section (Supplemental Note 10, Supplementary Figure 9). Further information, including comprehensive documentation of the SpatialData Python library, tutorials, example datasets, and a contributor guide are available online (<u>spatialdata.scverse.org</u>).

## Discussion

Here, we have presented SpatialData, a flexible, community standards-based framework to store, process and annotate data from virtually any spatial omics technology available to date. The ability to easily create common coordinate systems to align datasets is a critical cornerstone to establish common coordinate frameworks (CCFs). CCFs will unlock new analysis approaches that facilitate robust comparison and reuse of samples across studies. In conclusion, the flexibility and easily accessible solutions provided by the SpatialData framework enable new analysis possibilities and enhance the reproducibility of integrated spatial analysis.

As the uptake of SpatialData continues to grow, its utility will increase further. In the future, we aim to extend SpatialData interoperability to R/Bioconductor, provide support for multiscale point and polygon representations, and support cloud-based data access both programmatically and via the visualization tool Vitessce^17^. In summary, SpatialData fills the important need for an open and universal data framework for spatial omics.

## Contributions

L.M., G.P., K.A.Y., and I.V. designed and authored the spatialdata library. L.M., G.P., K.A.Y., I.V., and J.M. authored the spatialdata storage specification. G.P, L.M, M.T. and R.S. wrote napari-spatialdata. H.V. designed and prototyped the spatialdata-plot with input from T.T, G.P. and L.M., T.T, G.P. and H.V. implemented spatialdata-plot. E.H performed the analysis on the (Xenium & Visium) breast cancer dataset with input from L.M, G.P, and K.A.Y. O.S., F.J.T., and J.M. supervised the work.

## Data and code availability

SpatialData is available as a Python package via pip, and comes together with an extensive set of examples and tutorials, that can be accessed from scverse.org. Furthermore, researchers interested in contributing can join the discussion on the OME-NGFF specification and the SpatialData design directly on Github^18^. All scripts to reproduce the analysis can be downloaded from:

https://github.com/scverse/spatialdata-notebooks/tree/main/notebooks/paper_reproducibility.

## Acknowledgements

We would like to acknowledge the following individuals for their contributions: Danila Bredikhin for participating in a hackathon in Basel (April 2022) focused on discussions on representations for multiple modalities and in the scverse ecosystem; Wouter-Michiel Vierdag, Bishoy Wadie, Christian Tischer, Sebastian Gonzalez Tirado, and Leon Hetzel for attending a hackathon in Heidelberg (June 2022) and Ilia Kats for original prototype of spatial muon; Artem Lomakin for his contributions to discussions about alignment of clones and niches; Olga Lazareva for contributing work on clonality for the breast cancer study during the de.NBI BioHackathon SpaceHack project in Lutherstadt-Wittenberg (December 2022); organizers and participants of the de.NBI BioHackathon SpaceHack project. We appreciate the contributions from Helena L. Crowell, Constantin Ahlmann-Eltze, and Mike Smith for their prototype implementation of R readers for OME-Zarr and SpatialData objects as well as Laurens Lehner, Michal Klein, Wouter-Michiel Vierdag and Ilan Gold for their valuable contributions to spatialdata-io during a hackathon in Heidelberg (April 2023); Quentin Blampay for contributions to spatialdata-io; Artem Shmatko and Wouter-Michiel Vierdag for contributions to the implementation of the napari lasso tool; Matt McCormick for discussions around the usage of the packages SpatialImage and MultiscaleSpatialImage; Will Moore for discussions on OME-NGFF and technical support on OME-Zarr; John Bogovic for developing the OME-NGFF transformation specification; Joel Lüthi, Tong Li, Clarence Mah, Benjamin Rombaut and Lotte Polaris for discussions during the SpatialData meetings; Group members of Stegle and Theis lab for helpful discussions.

## Supplementary Information

### Supplementary Note 1: Challenges in spatial multi-omics data processing

#### Integration of spatial omics technologies for holistic views of tissue architecture

Spatial omics technologies are being applied to virtually all fields of biology, ranging from basic biological questions in model systems^19, 20^, the study of disease states in big cohorts of human patients^21, 22^, to the first clinical applications^23^. Current spatial omics technologies differ by readout (e.g., transcriptome, proteome, morphology), resolution and comprehensiveness^24^. For example, Visium profiling gives whole-transcriptome readouts at the cost of supercellular resolution: each Visium circular capture location describes the aggregated expression of up to dozens of cells^25^. On the other hand, single-molecule hybridization and in-situ sequencing technologies are capable of resolving millions of transcripts with diffraction-limited subcellular resolution, at the cost of a smaller panel of targets (usually up to a few hundreds)^26, 27^. Increasingly, multiple complementary technologies and profiling approaches are applied to the same samples^28^. However, these efforts are stymied by a lack of methods to integrate and jointly analyze different modalities^2, 4^.

#### Landscape of spatial omics analysis tools

There are several existing tools for analyzing spatial omics data, which provide complementary functionality. While some functionalities offered by SpatialData are also provided by existing solutions (Table S1), we identified 4 key outstanding challenges that are exclusively addressed by SpatialData:

- Support for large image data
- Spatial alignment of multimodal spatial omics data
- Cross-modality aggregation
- Interactive annotation

##### Integration of large image data

Images provide important complementary information to molecular profiles. For example, H&E images can provide critical histological and anatomical context for molecular measurements. Multidimensional images are generally stored as dense arrays which can be 10s-100s GB in size. Owing to their size, images require special consideration for performant processing and integration with molecular profiles. For example, lazy loading and multiscale representations only load the required portions and detail level of an image required for a given processing operation and thus allow images larger than the available memory to be processed. As a result, image processing software has largely been siloed from molecular profile analysis software. Thus, there is the need for a spatial omics analysis framework that fully embraces image data so image features can be integrated with molecular features.

##### Alignment of multimodal spatial omics data

Each spatial omics modality measures specific features of molecular architecture (e.g., RNA expression, morphology, chromatin accessibility). To build a holistic understanding of tissue architecture, it is beneficial to be able to combine complementary modalities and transfer quantifications and annotations between them. For example, if two regions of interest are identified by histological features in an H&E image, we may want to aggregate and compare the molecular profiles of protein abundance and RNA expression in those ROIs to study the signaling that gives rise to the distinct histological features. In order to achieve such an integration, disparate datasets need to be aligned into a common coordinate system (CCS). Computationally, this requires functionality to spatially transform all data types, which is non-trivial, as there are many types of data in spatial omics datasets (e.g., multidimensional arrays, polygons, points). Further, multiple aligned coordinate spaces may be required to represent the multiscale nature of biological architecture (e.g., an organ-specific coordinate system, and a tissue-specific coordinate system). Thus, to achieve practical multimodal spatial alignment, a rich transformation system with support for multiple coordinate systems is required.

##### Cross-modality integration

In addition to aligning multiple datasets, it is necessary to transfer spatial annotations between datasets in order to efficiently use complementary insights that are provided by each modality. In the context of the example above, transferring the spatial annotations of ROIs derived from the H&E image to the molecular profiles enables stratification and comparison of the signaling in distinct biological compartments. Such a comparison would not have been possible without transferring the annotations from the H&E image to the molecular profiles. While conceptually straight-forward, transferring spatial annotations across modalities is non-trivial to achieve in practice, as the corresponding aggregation operations must be applied to different data types (e.g., multidimensional arrays, polygons, points). Thus, to achieve cross-modality integration, a uniform interface for aggregating all data types present in spatial omics datasets is indispensable.

##### Interactive annotation

Interpreting spatial omics datasets requires input from domain experts. For example, it is often desirable to annotate regions of interest based on histological or anatomical features. Such annotations are necessary to compare distinct anatomical compartments and provide ground truth for training and validating analysis algorithms. Owing to the size and heterogeneity of spatial omics data types, interactive analysis of spatial omics datasets requires a performant viewer that can represent diverse data types.

##### Comparisons of storage and objects for spatial omics data handling

We evaluated the libraries in Table S1 on the basis described below.

##### Data types

Data types include the type of representations that constitute a building block of the spatial omics experiment:

-Raster images: multiplexed microscopy images.
-Raster labels: segmentation masks.
-Multiscale raster: pyramid-like representation of large microscopy images or labels.
-Polygons: list of polygons representing regions of interest (e.g. pathology annotation, tissue regions)
-Regular shapes: similar to polygons, used to represent capture locations of array-based technologies (e.g. circles for Visium, squares for DBiT-seq etc.)
-Points: list of dimensionless points, used to represent e.g. transcripts locations.
-Features matrix: gene or protein expression matrix.
-Annotation matrix: experiment metadata, cluster annotation etc.
-Graphs: neighbors graphs between regions (cells, spots, shapes) or features (genes, proteins).

##### Operations

These include operations to process spatial omics experiments, such as obtaining crops or slices of the data, summary statistics and data type conversion.

-Points aggregation: compute summary statistics between points and regions (labels, polygons or shapes) such as counting the number of transcripts across segmented cells.
-Geometry intersection: set operations between polygons, labels or shapes.
-Transforms: transform elements (images, regions and points) between coordinate systems
-Coordinate systems: support for specifying different coordinate systems for each element (e.g. pixel-based coordinate systems versus global physical coordinate systems).

##### Plotting

Plotting can be static or interactive.

**Table S1:**
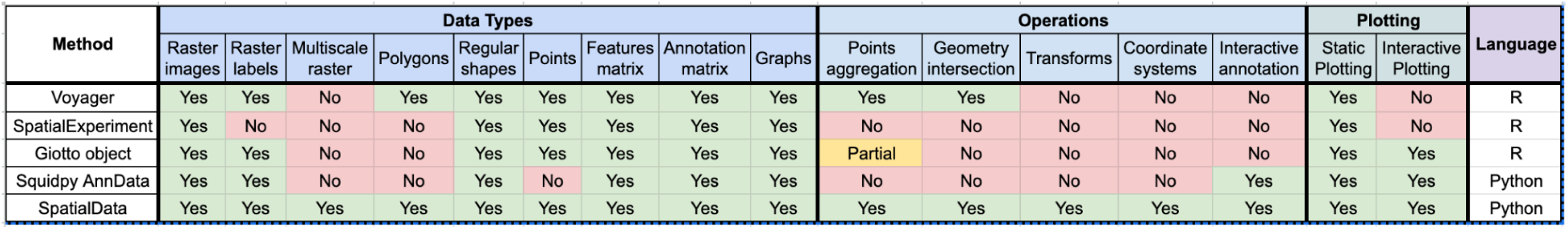
Existing spatial omics analysis lacks support for large images, aligning multiple datasets to a common coordinate framework, and interactive annotations.

### Supplementary Note 2: Universal representation of spatial omics data

Spatial omics datasets are challenging to load and integrate as they comprise different data types. The SpatialData storage format and in-memory representation supports 5 primitive SpatialElements to represent different datasets and data types: Images, Labels, Points, Shapes, and Tables (explained below in detail). These SpatialElements allow for representing the raw and derived data from a wide range of spatial omics assays, and we provide convenient reader functions for most common spatial omics data formats (Table S2). Multiple SpatialElements can be grouped together in a SpatialData object. SpatialData objects can represent a single dataset or multiple datasets. SpatialData objects are stored on disk in the SpatialData format, which is built upon the Zarr implementation of the OME-NGFF ^6, 7^ specification (Figure S1). OME-NGFF is a community-driven data standard with readers in Python, Java, and JavaScript, enhancing the interoperability of the SpatialData storage format. Using the standardized metadata from OME-NGFF also improves the accessibility and reproducibility of SpatialData.

-**Images**: images are raster data that store high-resolution microscopy images. They are stored as Zarr arrays and are represented in-memory as a (multiscale) *SpatialImage* class^29^. *SpatialImage* inherits from *xarray*^30^ and *xarray-datatree*^31^ for representing and manipulating high-dimensional arrays with named coordinates.
-**Labels**: labels are raster data that contain regions of interest such as segmentation masks. They are stored similarly to images as Zarr arrays on disk and represented in-memory as (multiscale) *SpatialImage*.
-**Shapes**: shapes are polygon data that contain regions of interest such as cell segmentations, capture locations of array-based spatial transcriptomics data or other types of ROIs. They are stored as a series of arrays that contain coordinates and offsets of the polygons as Zarr arrays on disk and represented in-memory as Shapely^32^ objects in GeoPandas^33^ dataframes.
-**Points**: points contain large collections (typically order of millions or billion) of coordinates and annotations such as transcripts locations and their associated metadata. They are stored as a parquet file on disk and represented in-memory as a lazy object with a DaskDataFrame^34^.
-**Tables**: tables store molecular profile information (gene expression, protein expression etc.) and associated metadata for observations and variables. It also stores the adjacency matrix of spatial graphs as well as any relevant additional metadata. It is stored on disk and represented in memory as AnnData^35^.

To illustrate how the SpatialData format works for standard spatial omics assays, we have converted 42 fields of view from 7 different technologies in the SpatialData format and made them available online (Table S3). We anticipate these standardized datasets will help analysis methods developers benchmark their methods across different spatial omics modalities.

**Supplementary Figure 1.**
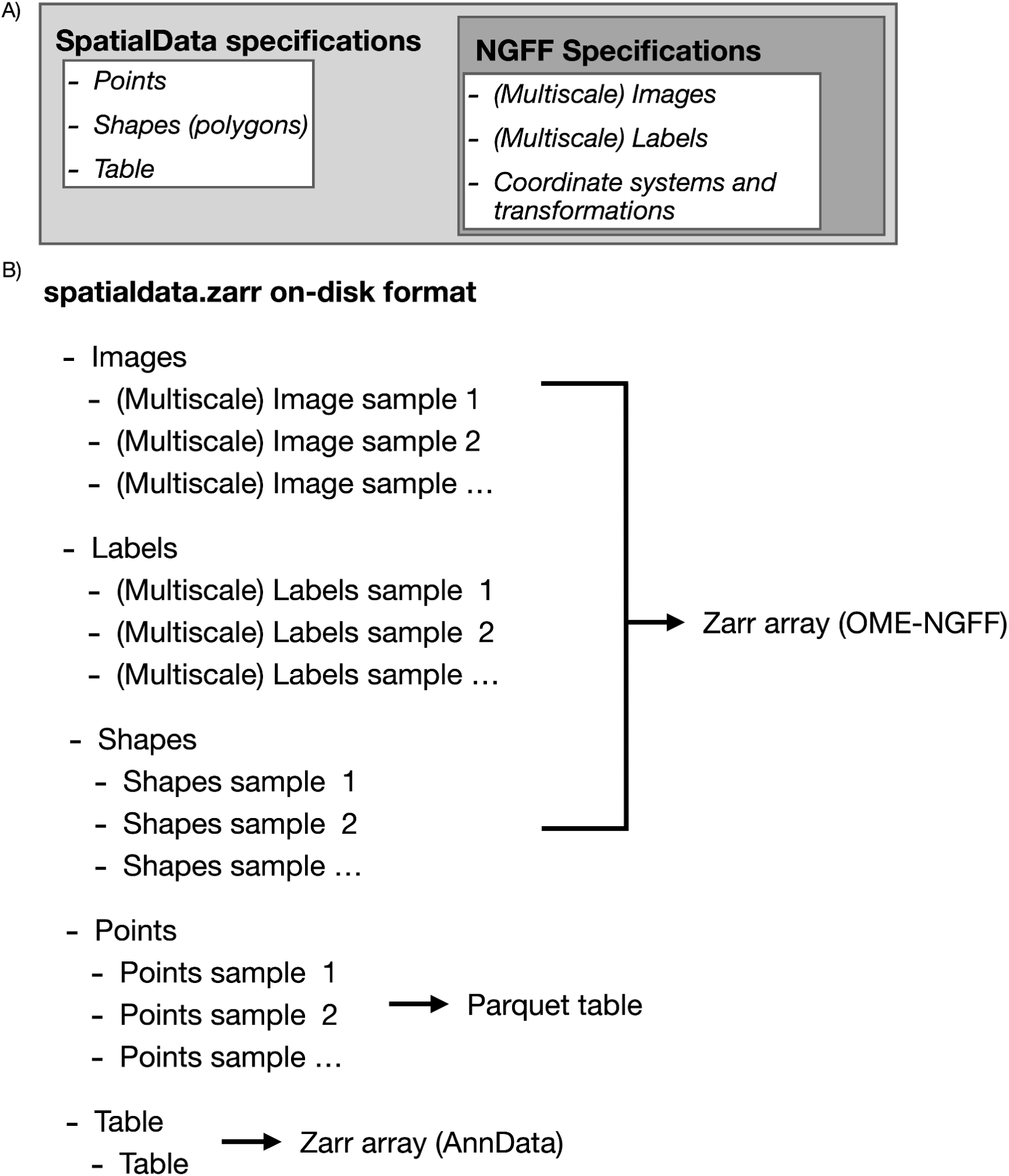
Schematic of the SpatialData storage layout in OME-Zarr. **(A)** The SpatialData storage format builds on the OME-NGFF specification by prototyping new data types required to store spatial omics information. Coordinate systems and transforms as well as tables are currently in review by the community. Points and shapes will be submitted to the community for review in a next phase to ensure interoperability. **(B)** Storage format of SpatialData: it consists of one Zarr container with nested folder structure, one for each SpatialData element. Each of the elements is saved as Zarr arrays, except for the Points, which are stored as an Apache Parquet file. We want to highlight that the Parquet file storage for points might change in the near future when a Zarr alternative will be implemented.

**Table S2.** | The SpatialData library provides reader functions for canonical spatial omics technologies and vendor-specific file formats. Shown are technologies, associated reader function in the SpatialData library and the set of SpatialData elements used to represent the data.

| Vendor/Technology | Reader function | Data | SpatialData elements |
| --- | --- | --- | --- |
| NanoString CosMx | cosmx | Transcripts locations | Points |
|  |  | Raster Images | Images |
|  |  | Segmentation masks | Labels |
|  |  | Gene expression | Table |
|  |  | Fluorescent marker intensity | Table |
|  |  | Metadata | Table |
| 10x Genomics Xenium | xenium | Transcripts locations | Points |
|  |  | Raster Images | Images |
|  |  | Cell segmentation | Shapes |
|  |  | Nuclei Segmentation | Shapes |
|  |  | Gene expression | Table |
|  |  | Metadata | Table |
| 10x Genomics Visium | visium | Raster Images | Images |
|  |  | Circular regions | Shapes |
|  |  | Gene expression | Table |
|  |  | Metadata | Table |
| CyCIF (MCMICRO output) | mcmicro | Raster Images | Images |
|  |  | Segmentation masks | Labels |
|  |  | Protein expression | Table |
|  |  | Metadata | Table |
| Imagine Mass Cytometry (Steinbock output) | steinbock | Raster Images | Images |
|  |  | Segmentation masks | Labels |
|  |  | Protein expression | Table |
|  |  | Metadata | Table |

**Table S3.** SpatialData provides a growing collection of sample datasets in the SpatialData storage format. We are continuing to add more datasets as they become available. For an up to date list, please see the online documentation: https://spatialdata.scverse.org/en/latest/tutorials/notebooks/datasets/README.html

|  | Tissue Type | Number of samples/sections | File size |
| --- | --- | --- | --- |
| NanoString CosMx | Non-small cell lung cancer (NSCLC)<sup>36</sup> | 30 adjacent slides from the same sample | ~4.2 GB |
| 10x Genomics Xenium | Breast cancer<sup>13</sup> | 2 samples with overlapping area | ~14.5 GB |
| 10x Genomics Visium | Breast cancer<sup>13</sup> | 1 sample | ~1.8 GB |
| CyCIF (MCMICRO output) | Breast cancer<sup>37</sup> | 1 sample | ~250 MB |
| MERFISH (Allen Institute prototype pipeline) | Mouse brain<sup>38</sup> | 1 sample | ~50 MB |
| MIBI-TOF | Colorectal cancer (CRC)<sup>39</sup> | 3 slides | ~25 MB |
| Imaging Mass Cytometry (Steinbock output) | Patient 1: SCCHN (head and neck cancer)<br>Patient 2: BCC (breast cancer)<br>Patient 3: NSCLC<br>Patient 4: CRC<sup>40,41</sup> | 4 patients, 14 images | ~820 MB |

### Supplementary Note 3: Spatial query interface

When operating on large data, it is helpful to be able to subdivide the dataset into multiple regions. This can be to extract specific anatomical regions or to divide the dataset for parallel processing. Via the spatial query interface, users can request the data contained in a bounding box in a specific coordinate system. The result of this subset operation is returned as a SpatialData object. In future work, we will extend the coordinate system for more advanced queries such as compound queries on both spatial coordinates and feature values (e.g., all cells within region X and belonging to cluster Y). To help analysts learn how to use the spatial query functionality, we have created a tutorial that can be downloaded as a Jupyter notebook available online via the SpatialData online documentation (Figure S2).

**Supplementary Figure 2.**
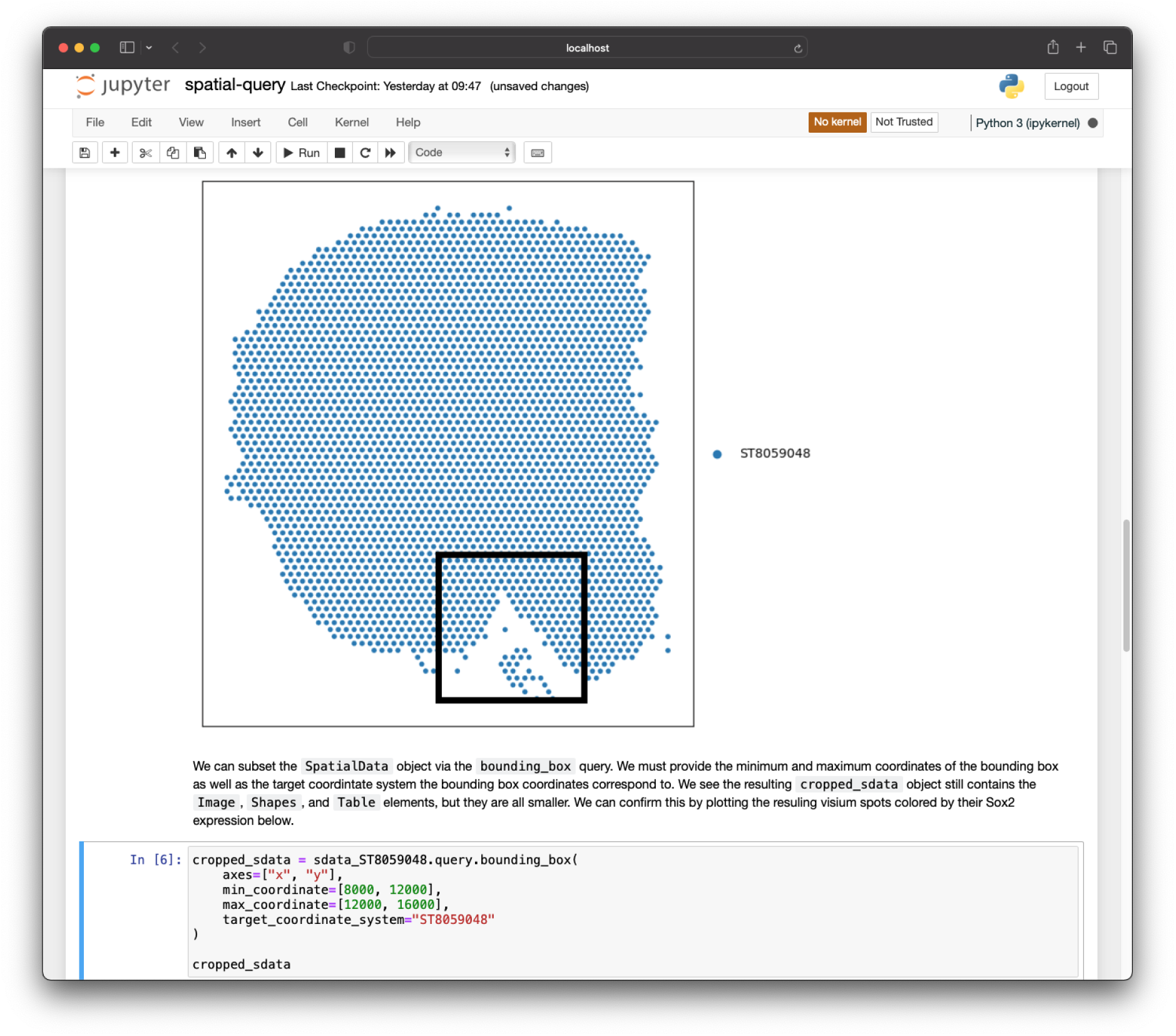
Illustration of the SpatialData query function. Shown is a screenshot for the spatial query tutorial from the online SpatialData documentation. The tutorial illustrates the selection of data by spatial bounding boxes defined in the common coordinate system. The full example can be found here in the “spatial query” notebook in the online documentation (<u>link to tutorial</u>).

### Supplementary Note 4: Aggregation operations operate interchangeably between data representation

Aggregation operations form the foundation for transferring quantifications and annotations across modalities in multimodal analyses. The SpatialData framework enables the aggregation (also referred to as accumulation in image processing) of data stored in any SpatialElement into any set of target geometries or masks (Figure S3). For instance, it is possible to count the number of single molecules for a specific gene within polygon geometries representing cells. Similarly, those same molecules could be counted within image masks representing the cytoplasm of the cells. Another example is averaging cell gene expression within a given anatomical region.

SpatialData offers the flexibility to apply standard aggregation operators to the data (count, sum, mean, standard deviation) to any SpatialElement. Additionally, SpatialData provides an interface for applying aggregation operators defined by the user. Leveraging the SpatialData common coordinate system, aggregations can be performed between two SpatialElements that have different spatial scale and/or do not fully overlap. To help analysts learn how to use the aggregation system, we have made a tutorial available in the SpatialData online documentation (<u>link to tutorial</u>).

**Supplementary Figure 3.**
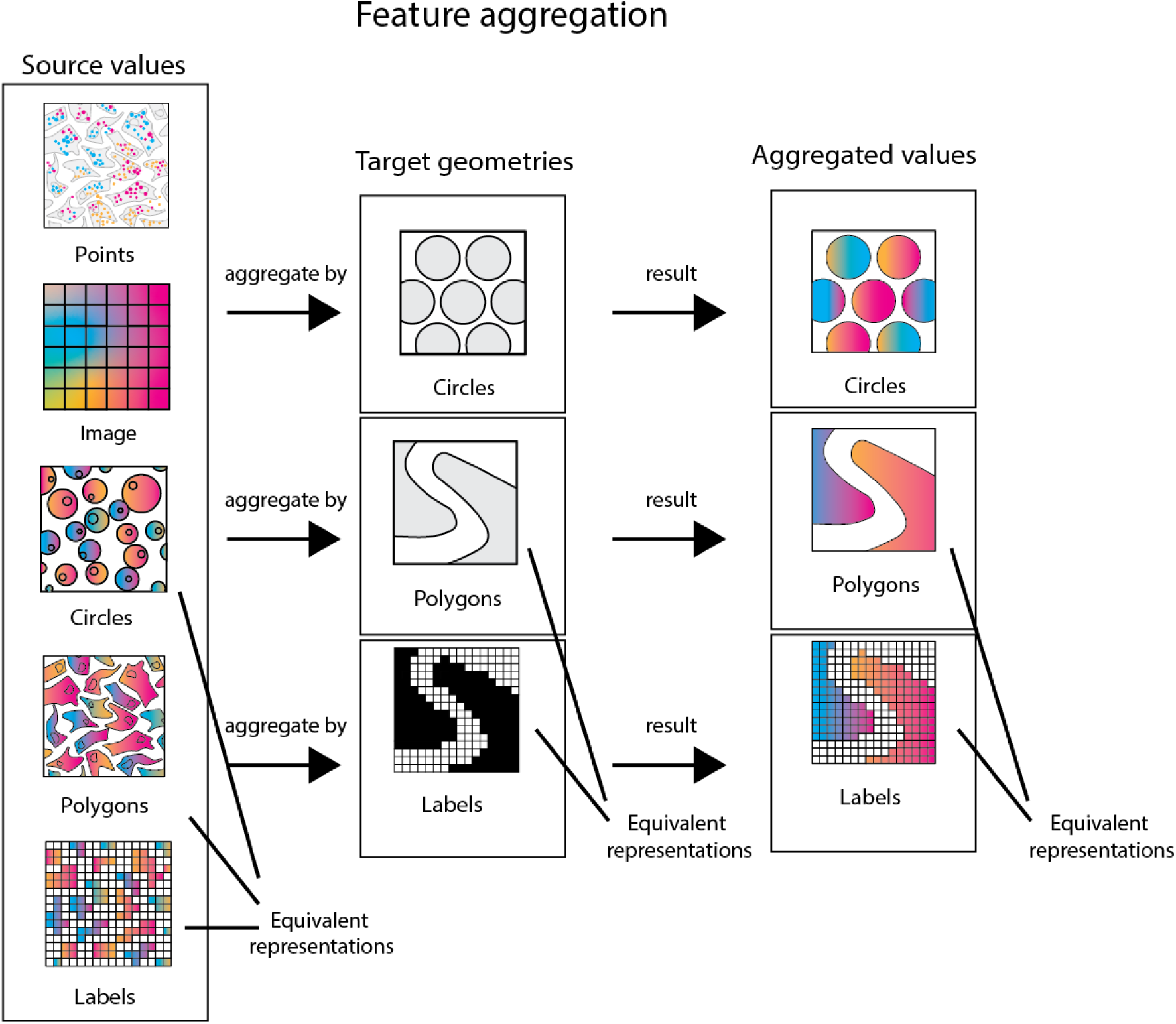
Schematic representation of the SpatialData aggregation operations. SpatialData provides a uniform programmatic interface for aggregating observations of any spatial omics modality by spatial annotations. Aggregation operations take values from a source SpatialElement and spatial annotations from a target SpatialElement as inputs. The aggregation operation groups the input values by the target geometries, applies an aggregation function (e.g., mean, sum, count) to the grouped input values and returns the aggregated values. Owing to the storage interface of SpatialData, aggregations can be applied in a uniform manner to all types of spatial omics data.

### Supplementary Note 5: napari-spatialdata for interactive visualization and annotation of spatial omics data

**Supplementary Figure 4.**
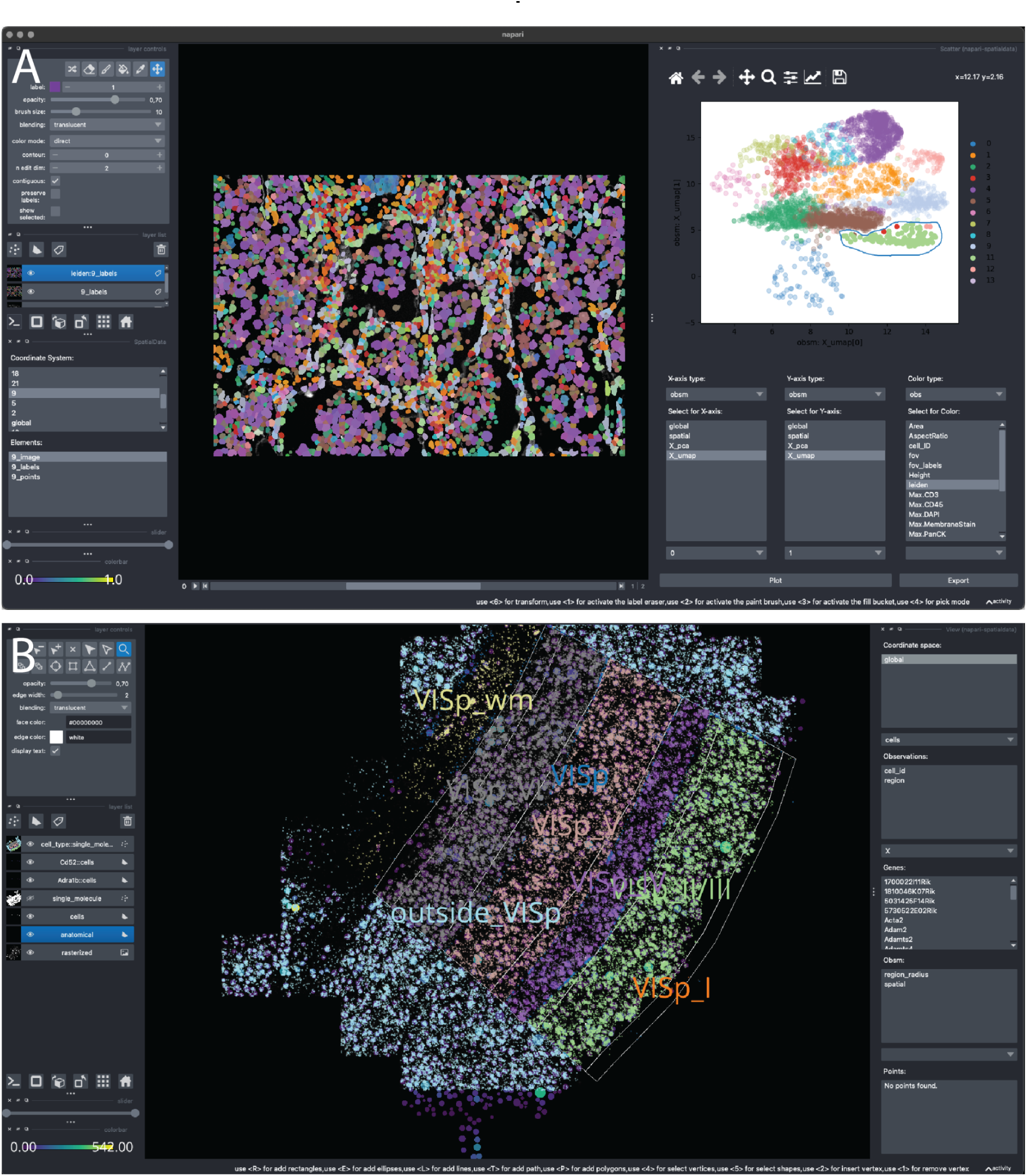
Example of using napari-spatialdata to visualize and annotate various datasets. Napari-spatialdata interactively visualizes any of the SpatialElements (images, labels, points, shapes) with their associated annotations (such as gene expression, cluster annotations etc.). Embeddings of molecular profiles (e.g., t-SNE, UMAP) can be interactively explored via the scatter plot widget. Spatial annotations can be made via drawing of regions in the napari viewer. The spatial annotations are exported in the SpatialData object so that they can be used in downstream analysis. **A.** NanoString CosMx dataset and interactive selection with a lasso from the UMAP plot computed from the cell gene expression and colored by Leiden clusters. The lasso tool in the scatterplot windows is being used to annotate a set of cells. The annotation can be visualized in space and can be exported for downstream usage. **B.** MERFISH mouse brain dataset (Allen Institute prototype MERFISH pipeline^38^) featuring gene expression, polygonal ROIs annotating anatomical regions and cell types assigned to single molecule points.

### Supplementary Note 6: Static plotting with spatialdata-plot

**Supplementary Figure 5.**
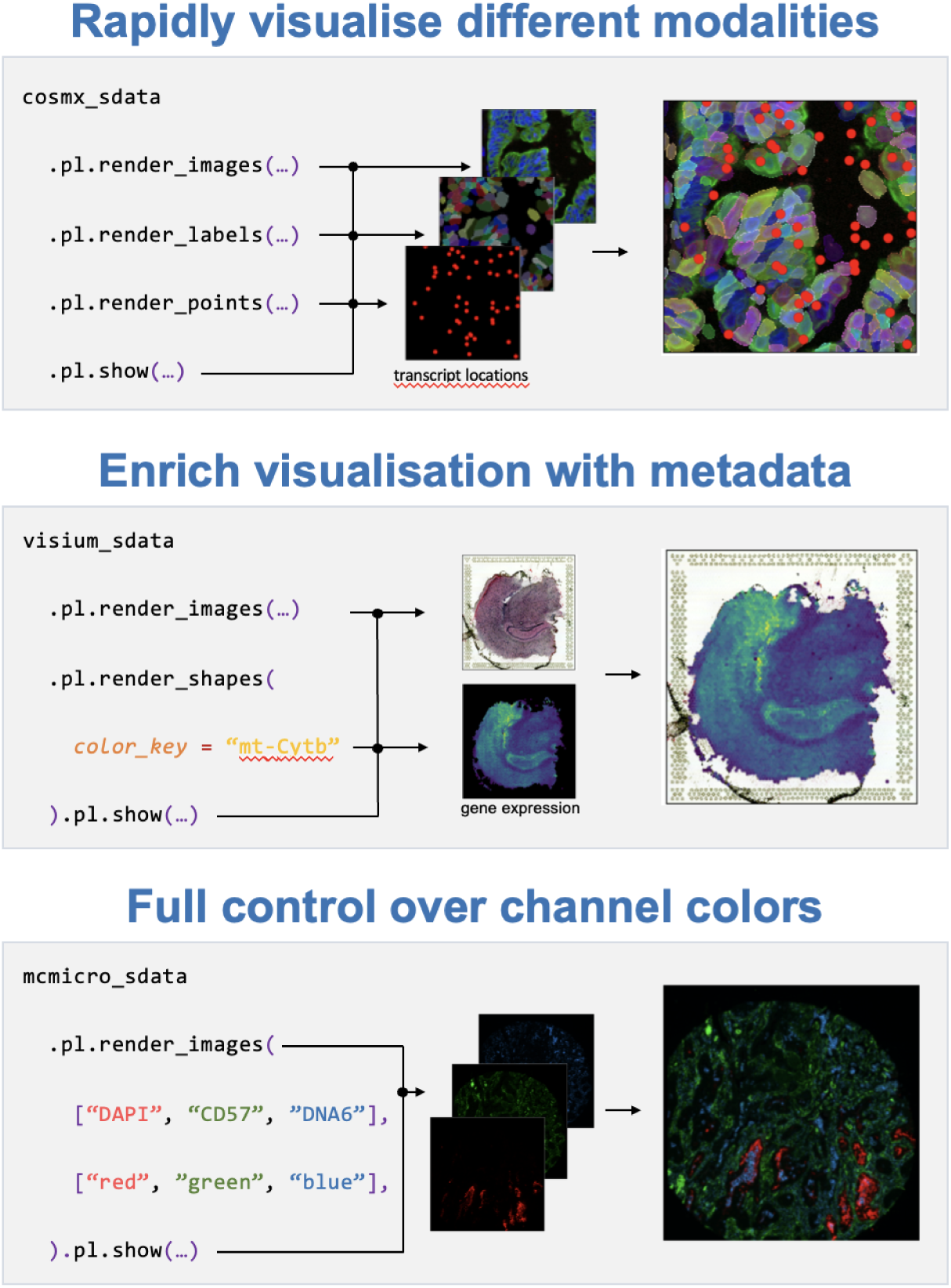
Illustration of functionality of the static plotting library spatialdata-plot. The library spatialdata-plot enables the streamlined visualization of complex multi-modality data. SpatialData’s plotting capabilities can be used to visualize different modalities. The user can specify which elements should be rendered (images, labels, points, shapes), and specify parameters for each plotted element. For example, shapes representing cells can be colored by a gene’s expression. The plotting library automatically accounts for transformations and alignments of the underlying common coordinate system,

### Supplementary Note 7: Training of deep learning models directly from SpatialData datasets

Deep learning algorithms are promising for multimodal integration and prediction. However, they require careful curation and preparation of training datasets. Dataset preparation is especially time consuming for spatial omics data because of the heterogenous data types, large dataset sizes, and the need for spatially-aligning multiple modalities. Leveraging the spatial query API, SpatialData comes with data loaders that are derived from the PyTorch Dataset class^42^, thereby facilitating data ingestion for deep learning applications (Figure 1D, Figure S6). The Dataset implementation uses the spatial query functionality to generate tiles from a SpatialData object. This implementation enables users to integrate SpatialData datasets with the rich Python deep learning ecosystem including models and infrastructure from MONAI.

A tutorial on how to use the PyTorch dataset loader is available as part of the online documentation (<u>link to tutorial</u>), where we use the SpatialData PyTorch dataset interface to train a MONAI DenseNet encoder on the breast cancer study discussed in the main text (Figure 2B). The tutorial demonstrates how to generate image tiles from an H&E image that is spatially aligned to one of the two Xenium datasets, by querying the image tiles around each Xenium cell using the SpatialData PyTorch Dataset class. We then use these data to train DenseNet to predict the cell type from each image tile.

**Supplementary Figure 6.**
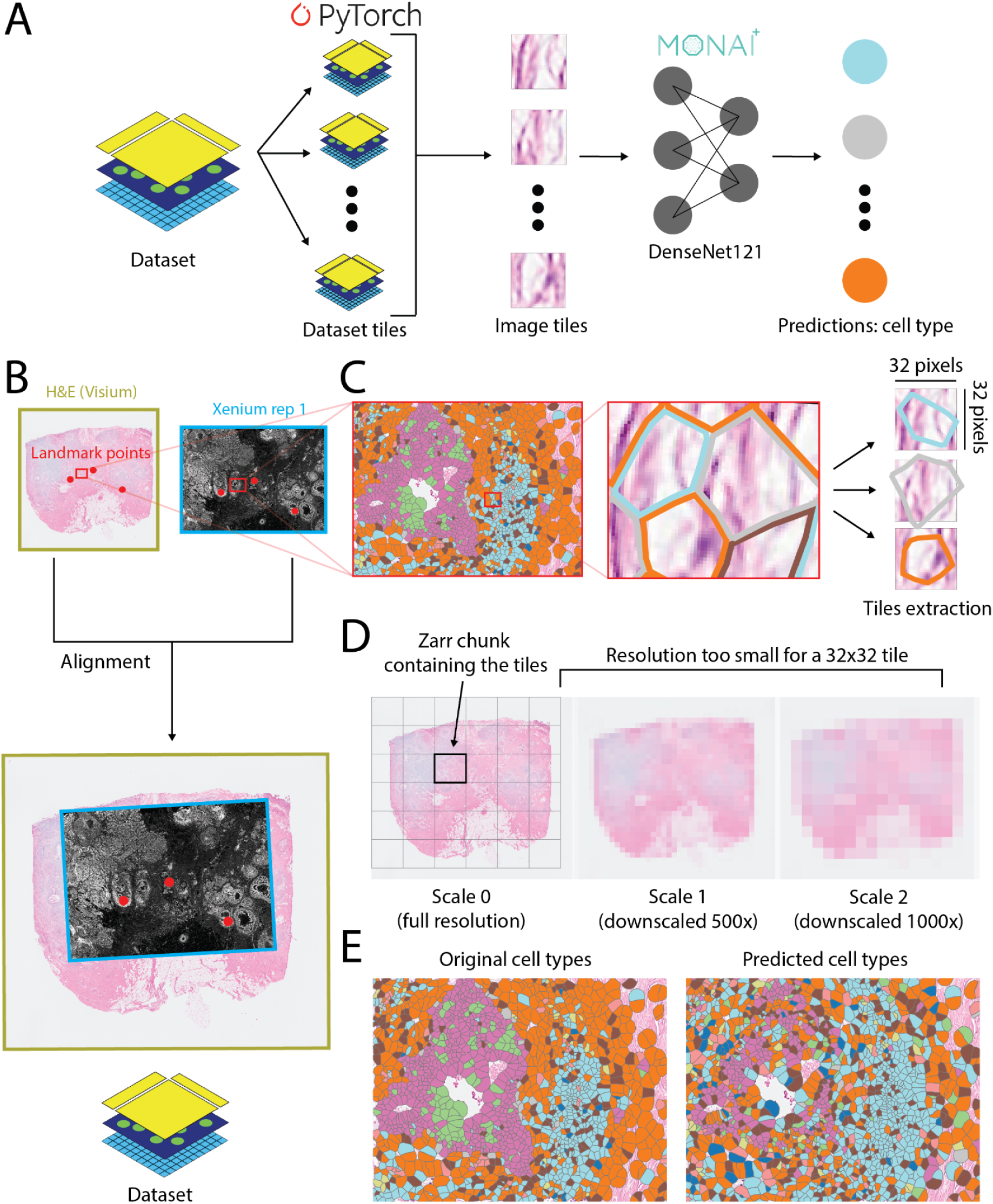
SpatialData facilitates the preparation of datasets for deep learning applications and it integrates with existing deep learning ecosystems. **(A)** SpatialData allows to generate PyTorch datasets that subset the original SpatialData object around desired locations. The resulting dataset tiles are valid SpatialData objects themselves. Here we construct image tiles around cells and we use them to predict the corresponding cell types using a DenseNet encoder. The model is provided by the MONAI framework, and this example shows how we can readily interface with existing deep learning ecosystems. **(B)** The generation of deep learning datasets harness the use of common coordinate systems to combine different spatially aligned elements, even in presence of diverse resolutions or affine transformations. Here are shown the H&E image and Xenium replicate 1 aligned datasets precedently introduced in Figure 2A. **(C)** Enlarged view of a portion of the two datasets, overlaying the cells from Xenium, colored by cell type, to the H&E image from Xenium. SpatialData allows to extract image tiles of the desired resolution (here 32×32 pixels) around the Xenium cells. **(D)** The tiling extraction process takes advantage of the multiscale representation and the chunked Zarr storage for efficient memory usage. The first allows the extraction of the tiles from the appropriate (downscaled) resolution, the second ensures that only the data chunk(s) containing the information about the tiles are loaded from disk. Note: the 500x and 1000x downscaling factors and the size of the chunks have been chosen for graphical purposes. Usually the downscaling factors and the chunk size are smaller. **(E)** Visualization of cell types predictions from the model. Note: since our focus in this example is to demonstrate the infrastructure, the network has been trained only for a few epochs and without optimizing the hyperparameters. This is reflected in the suboptimal accuracy of the predictions.

### Supplementary Note 8: Integration with squidpy

SpatialData is compatible with the scverse ecosystem ^12^. For example, a spatialdata can be integrated with squidpy functionalities, to compute various types of spatial summary statistics. We showcase how such integration works in a notebook in the online documentation (Figure S6).

**Supplementary Figure 7:**
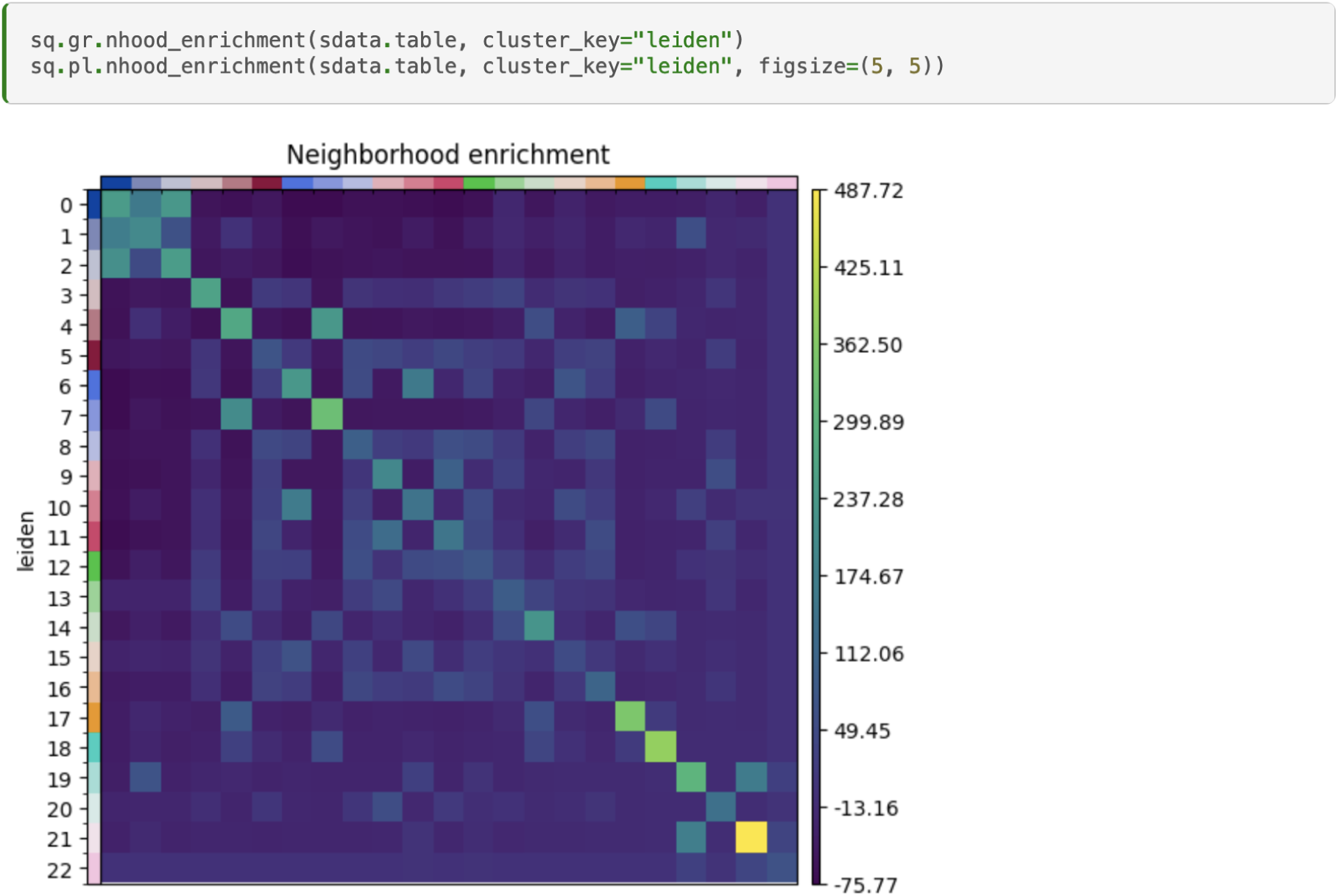
By building upon standard scientific Python data types, SpatialData can be readily integrated with existing packages such as squidpy. Shown is the result from a spatial neighborhood enrichment analysis on a 10x Genomics Xenium dataset. The rows and columns of the heatmap correspond to a cluster identified in the dataset, and each entry in the heatmap represent the enrichment score: a high enrichment score means that the two clusters are found to be enriched in spatial coordinates, i.e.,they are neighbors, while a low enrichment score means that the two cluster are not found to be neighbors across the tissue. See the “squidpy integration” example notebook in the online notebook for details <u>link to tutorial</u>.

### Supplementary Note 9: SpatialData operations for aggregating multi-modal datasets facilitates technological and methodological benchmarking

**Supplementary Figure 8.**
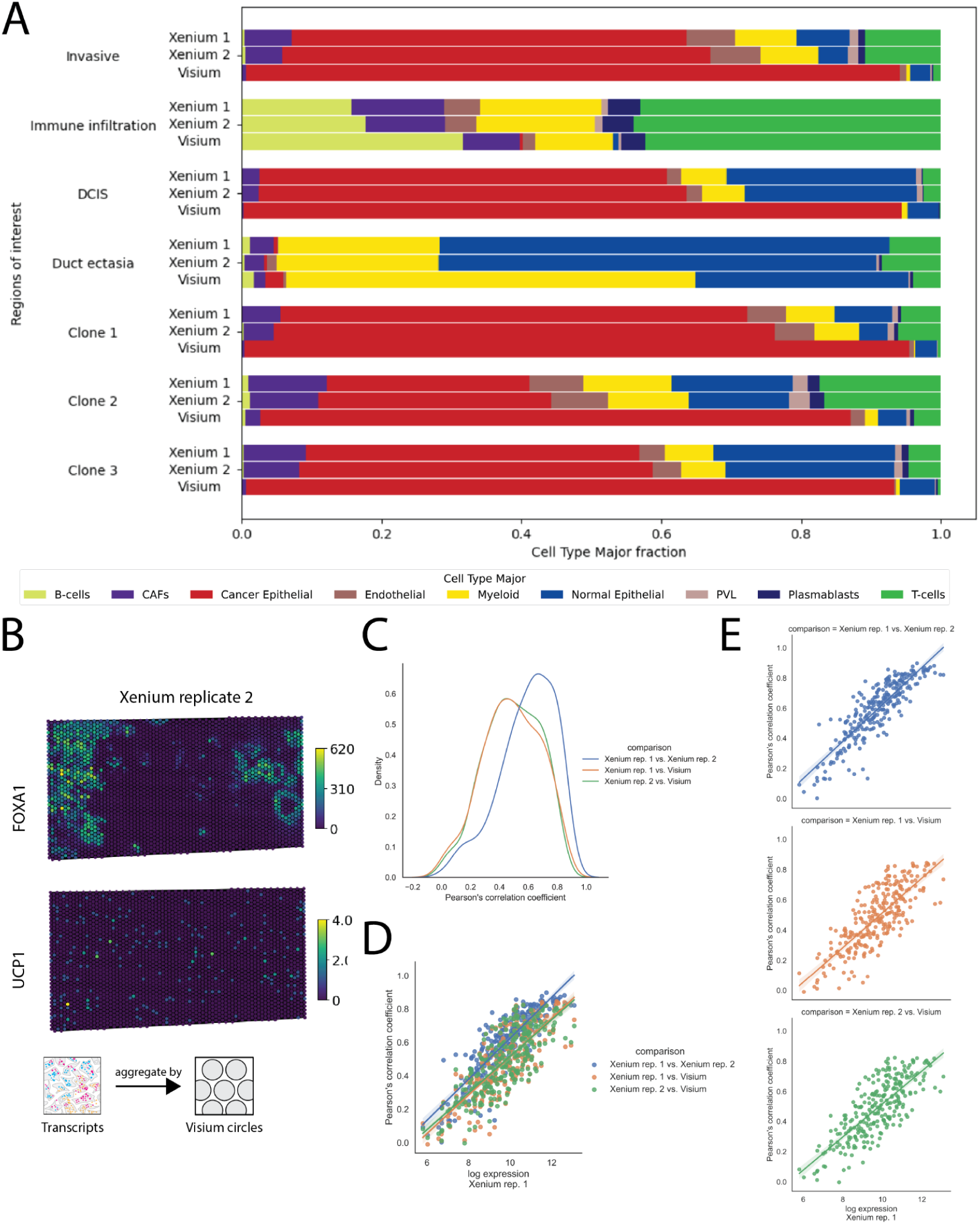
Supplementary results for the analysis presented in main text Figure 2. **(A)** Supplement of Figure 2D for the remaining ROIs and clones. Cell type proportions are computed over annotated regions of interests (ROIs) as well as clones across the Xenium replicates as well as the Visium dataset. For Visium the inferred cell type proportions using cell2location is shown. ROIs are selected based on histological features via the napari-spatialdata plug-in on the Visium-associated H&E image. The clones are inferred on the Visium data using CopyKat ^14^. **(B)** Supplement of Figure 2E for replicate 2. Aggregated transcripts for FOXA1 and UCP1 for Xenium replicate 2 over the Visium locations. **(C)** Supplement of Figure 2E. Overall density of Pearson’s correlation coefficient via pairs of sections is computed for each gene. **(D, E)** The relationship between the Pearson’s correlation coefficient between pairs of sections and overall gene expression is shown. Each point represents a gene.

### Supplementary Note 10: Representation of dataset with multiple overlapping field-of-views and different modalities

**Supplementary Figure 9.**
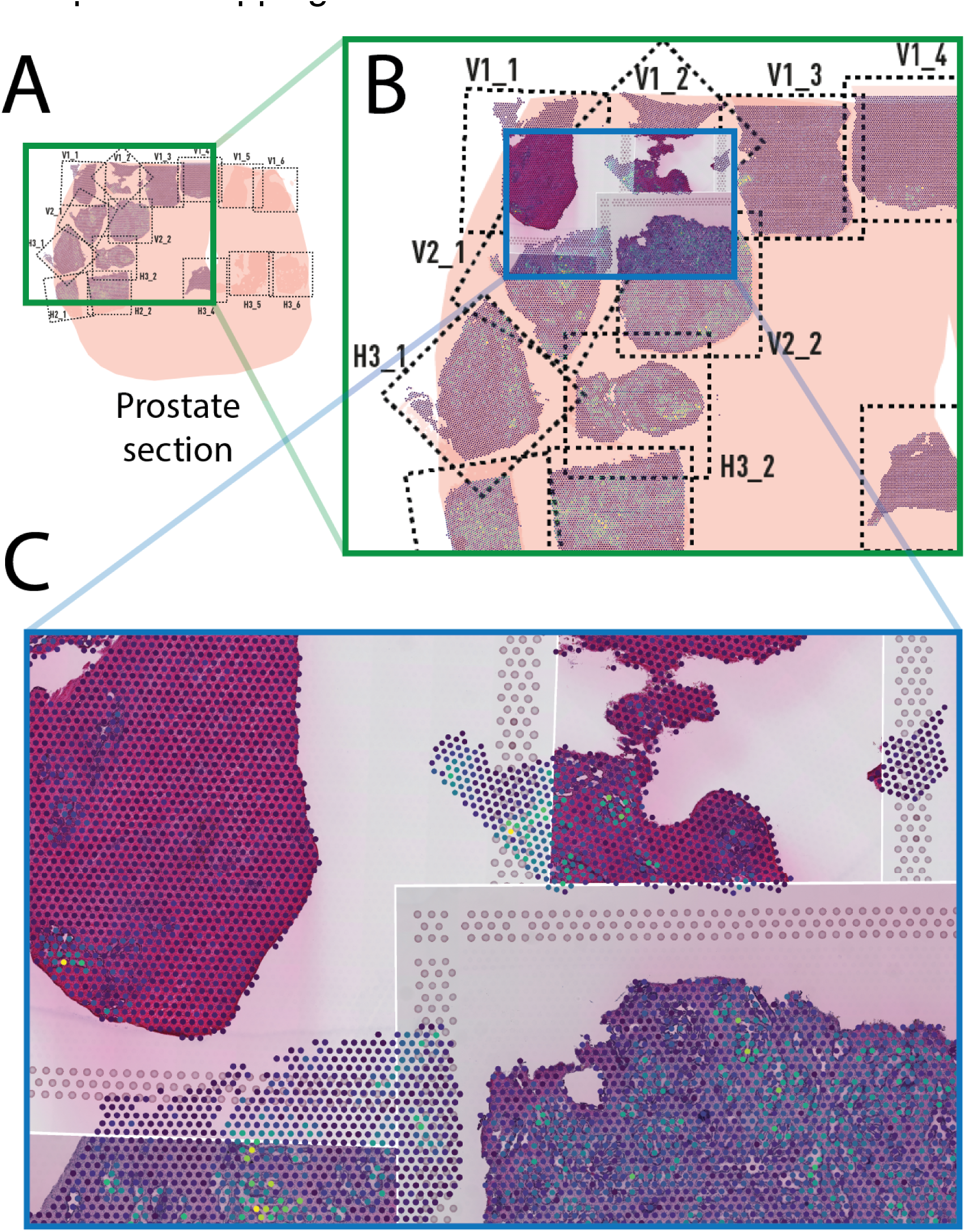
Example of using SpatialData to combine multiple datasets from a prostate cancer study. Shown is a common coordinate system constructed using data from Erikson *et al.* ^43^. The study comprises multiple Visium H&E and Spatial Transcriptomics^43^ datasets from multiple tissue samples, with partially overlapping fields-of-view distributed across the tissues. **(A)** Layout of the 15 fields-of-view for the Visium experiments for one of the tissues. The coordinate transformations as shown were derived using SpatialData (landmark-based alignment) mapping the data to global layout images shown in the original publication. **(B)** Screenshot of the visualization of all Visium datasets for one of the tissue samples in the context of the whole tissue coordinate system using napari-spatialdata. **(C)** Leveraging the SpatialData multiscale image representation and napari-spatialdata, we can view and explore all of the large images (15 images, ≈ 580 megapixels each) aligned together with the spatial gene expression. We can also visualize multiple modalities together, such as adding to the view also the Spatial Transcriptomics data. *The layout image used in the background in panels A and B has been made available in the original publication*^43^ *under the Creative Commons Attribution 4.0 International License. To view a copy of this license, visit* http://creativecommons.org/licenses/by/4.0/.

### Supplementary Note 11: Interactive visualization of a three dataset breast cancer study

**Supplementary Figure 10.**
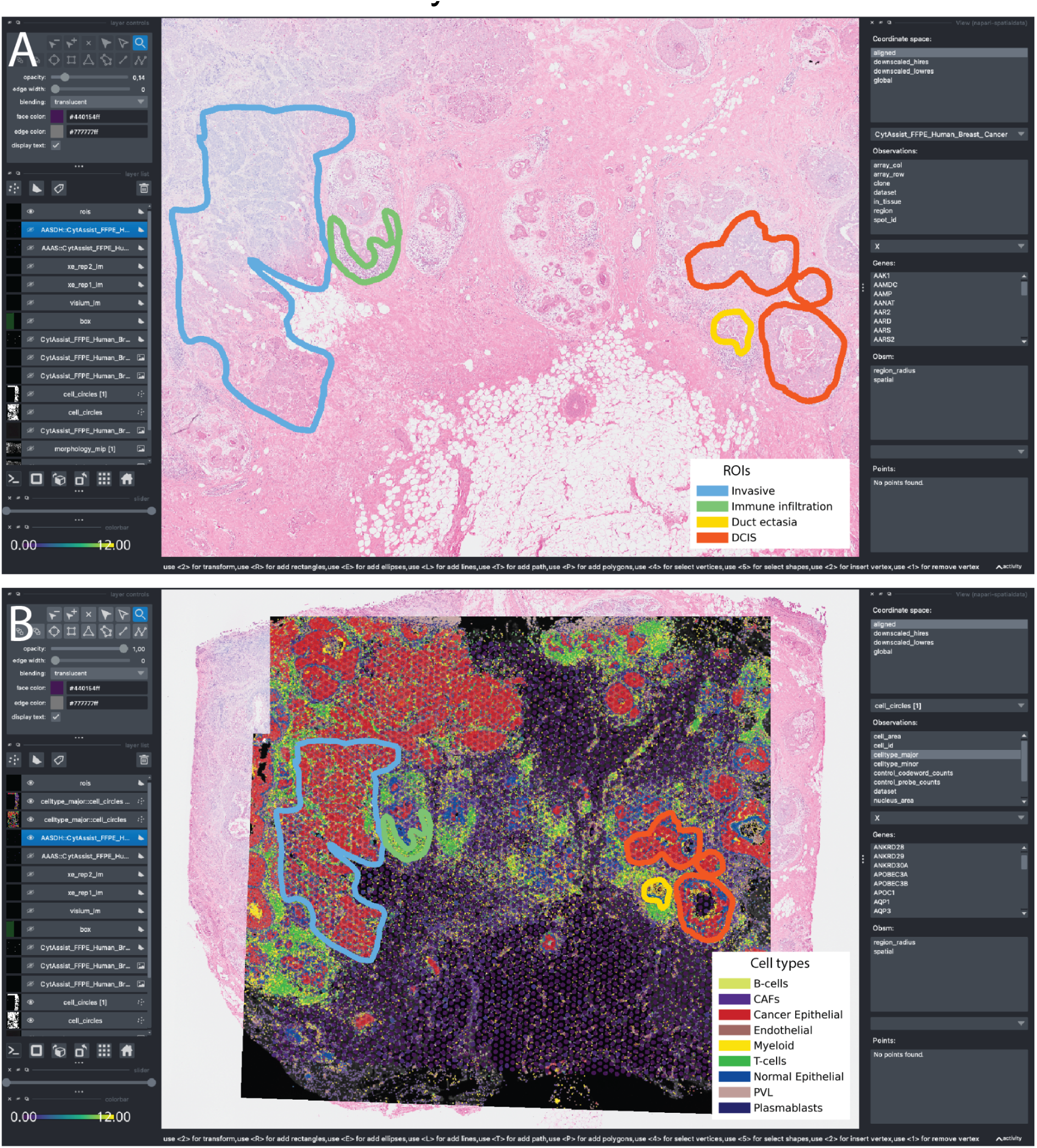
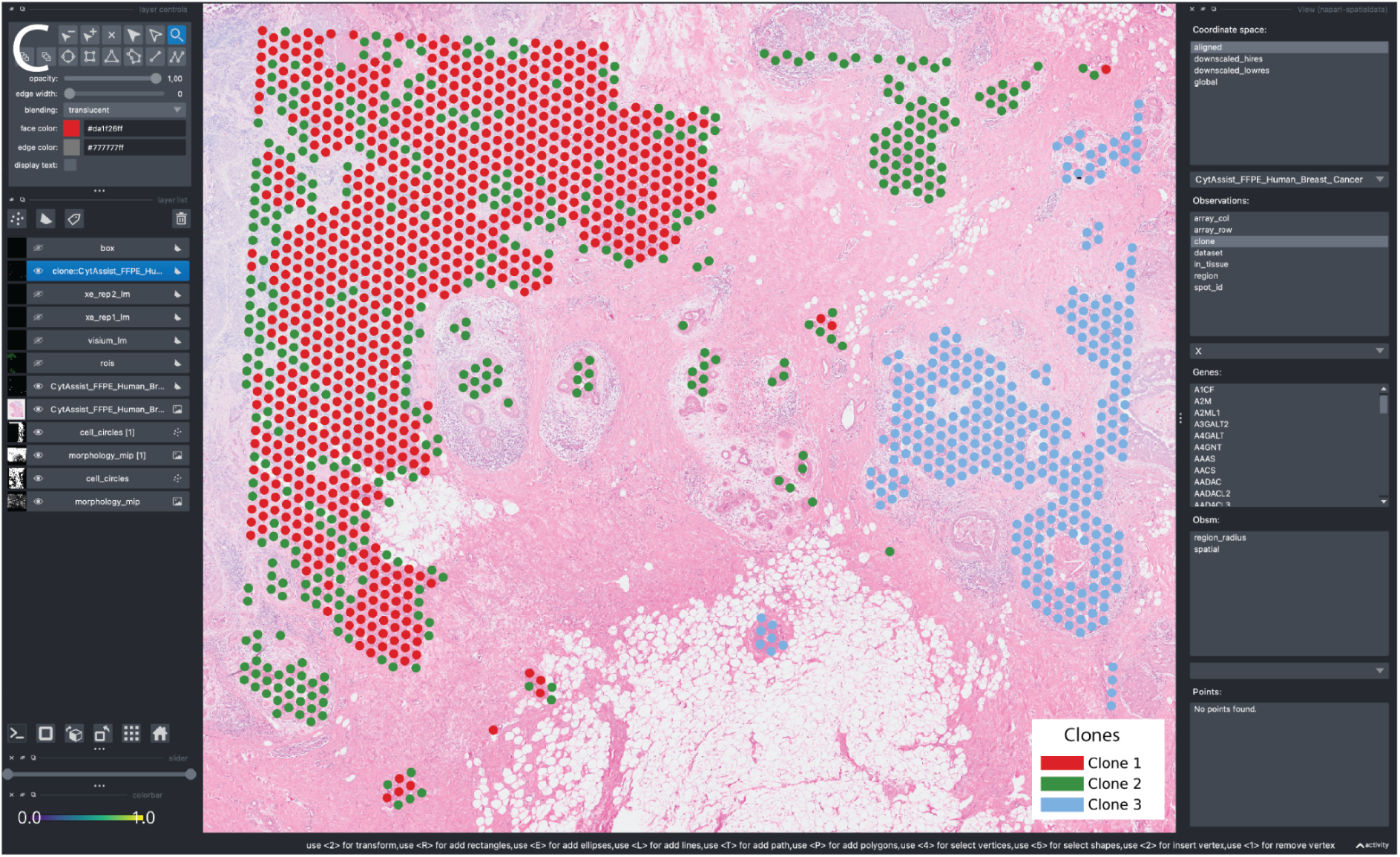
Visualization with napari of the Visium and the two Xenium datasets from the breast cancer study discussed in the main text (**Figure 2B**). A. Showing the H&E image from the Visium dataset annotated with the ROIs for anatomically relevant tissue compartments. B. Multimodal visualization overlaying the H&E image from the Visium data, the two immunofluorescence images associated with the Xenium data, the Visium array capture locations colored by gene expression (showed with transparency), the Xenium cells showing cell types and the four manually annotated ROIs. C. Visualization of the clones inferred from the Visium expression data.

## Methods

### SpatialData framework

The SpatialData framework comprises a core package and associated satellite packages napari-spatialdata, spatialdata-io and spatialdata-plot, compatible with Python 3.9 and above. All code is available on GitHub as part of the scverse organization, and is licensed under the permissive “BSD 3-Clause License’’. The project structures inherit from the scverse cookiecutter, and the napari plugin cookiecutter, thus implementing unit tests and pre-commit checks in a continuous integration setting. The documentation is built using Sphinx and hosted on Read the Docs. It includes API descriptions, example notebooks and a table with links to downloadable spatial omics datasets. Each dataset can be both downloaded in full (.zip), or even directly accessed from the cloud (public S3 storage).

We also provide a contribution guide and a technical design document to encourage adoption. Users can reach out to the core development team via the GitHub Issues bug tracking system. To encourage collaboration between the imaging and scverse communities, we have created a public chat stream in the imagesc Zulip messaging platform: https://imagesc.zulipchat.com/#narrow/stream/329057-scverse.

### Raw human breast cancer Xenium and Visium data

We downloaded the raw data from:

https://www.10xgenomics.com/products/xenium-in-situ/preview-dataset-human-breast.

### Loading Xenium and Visium datasets into SpatialData

The 10x Xenium and Visium readers from spatialdata-io were used to read in the data into SpatialData objects. For the Xenium datasets, the DAPI channel was stored as a multiscale Image, cell and nuclei segmentation masks and boundaries were stored as Shapes elements whereas the transcripts were stored as Points. The metadata and count matrices were stored as a Table in the SpatialData object. For the Visium dataset, the H&E image was stored as a multiscale Image, the array capture areas (circles) were stored as Shapes, and the count matrix and annotations were stored in the Table.

### Cell type annotation of the Xenium replicates

We annotated cells from the Xenium replicates using a publicly available scRNA-seq breast cancer atlas ^15^, comprising 9 malignant and normal cell types and 29 subtypes. After subsetting the atlas to the subset of 313 genes present in the Xenium panel, we applied the ingest method for label transfer as implemented in the Scanpy package (v1.9) ^9^ to annotate cells from the Xenium replicates. We transferred major cell type labels first (coarse-grained) and then within each class we mapped the minor cell types (fine-grained). In the current analysis only major cell types are shown. The 9 major cell types are B-cells, Cancer Associated Fibroblasts (CAFs), Cancer Epithelial, Endothelial, Normal Epithelial, Plasmablasts, and Pre-vasculature cells (PVL), and T-cells.

### Alignment to create common coordinate systems

We selected 3 landmark points from the images from the two Xenium replicates as well as the Visium dataset. The landmark points are to be selected on each of the images in the same order and there should be a 1-to-1 spatial correspondence between the sets of points. Xenium replicate 1 was used as the reference to which Xenium replicate 2 and Visium were aligned using the SpatialData function align_elements_using_landmarks. We used napari-spatialdata to annotate the landmark points and to view the result of the alignments. Internally, Dask’s lazy-loading and Zarr’s multiscale representation made it possible to performantly explore and zoom the datasets even in a low memory device like a standard laptop.

### Cell type interpolation for Visium

After alignment, the shared area between each cell, from the Xenium replicates, and the Visium locations, is computed. Then the cell type fractions are computed for each Visium location based on the surface fractions of the locations covered by each cell type. This is done for Xenium replicates 1 and 2 separately.

### Cell type deconvolution using cell2location

We used cell2location (v0.1.3) ^16^ to estimate the cell type fractions at the Visium locations. We used the aforementioned breast cancer atlas as the reference. For this task we operated on the subset of 313 genes present in the Xenium replicates and subset the Visium dataset and the breast cancer atlas to those genes. We set the default parameters as suggested in the cell2location tutorial https://cell2location.readthedocs.io/en/latest/notebooks/cell2location_tutorial.html. The analysis can be found at https://github.com/scverse/spatialdata-notebooks/tree/main/notebooks/paper_reproducibility. For visualization only cell types that contribute at least 5% per spot are taken into account. Then, each spot is normalized to have a total sum of 1.

### ROI selection with napari-spatialdata

After alignment four ROIs were selected based on the H&E image from the Visium dataset using the napari-spatial data plugin. These ROIs were then added to the aligned Xenium replicates as well. Each of the ROIs was selected based on their distinct micro-anatomical characteristics and then was labeled manually based on the underlying cell-type composition from the Xenium replicates.

### Clone detection on Visium using CopyKat

We used CopyKat (v1.1.0) ^4^ with the default parameters to estimate copy number states from the Visium count matrix. Then we performed hierarchical clustering, which identified 3 major clusters on the locations labeled as “aneuploid”. These three clusters were used as genetic subclones. We also transferred the clone labels to the overlapping cells from Xenium replicates. The clone labels were stored as a SpatialData table element. This analysis was conducted in R separately (see

https://github.com/scverse/spatialdata-notebooks/tree/main/notebooks/paper_reproducibility).

The Visium’s anndata table was saved in .h5ad format to be loaded and analyzed in R. The clone labels were then transferred back to SpatialData via .h5ad. There are ongoing efforts in the Bioconductor community to enable direct loading of anndata tables into R from Zarr, which would obviate the need for exporting as .h5ad (HDF5 format) when completed.

### ROI cell type fractions

We next computed, for each ROI and for each clone, the fractions of cell-types for the cells contained within them. The SpatialData aggregation APIs offer a convenient interface to compute these metrics, independently if what is being aggregated is a set of circles or polygons, and if the target region is a polygonal ROI or a set of circles defining a particular clone.

### Transcript aggregations

For each Visium spot we aggregated the transcripts from the Xenium replicates falling into each Visium spot. We performed this analysis for Xenium replicates 1 and 2 separately. This yields two aggregated count matrices that were saved as separate layers in the Visium’s SpatialData object’s table.

## References

1. Asp, M., Bergenstråhle, J. & Lundeberg, J. Spatially Resolved Transcriptomes-Next Generation Tools for Tissue Exploration. Bioessays e1900221 (2020) doi:10.1002/bies.201900221.

2. Rao, A., Barkley, D., França, G. S. & Yanai, I. Exploring tissue architecture using spatial transcriptomics. Nature 596, 211–220 (2021).

3. Rood, J. E. et al. Toward a Common Coordinate Framework for the Human Body. Cell 179, 1455–1467 (2019).

4. Palla, G., Fischer, D. S., Regev, A. & Theis, F. J. Spatial components of molecular tissue biology. Nat. Biotechnol. (2022) doi:10.1038/s41587-021-01182-1.

5. Wilkinson, M. D. et al. The FAIR Guiding Principles for scientific data management and stewardship. Sci Data 3, 160018 (2016).

6. Moore, J., et al. OME-Zarr: a cloud-optimized bioimaging file format with international community support. bioRxiv 2023.02.17.528834 (2023) doi:10.1101/2023.02.17.528834.

7. Moore, J. et al. OME-NGFF: a next-generation file format for expanding bioimaging data-access strategies. Nat. Methods 18, 1496–1498 (2021).

8. Palla, G. et al. Squidpy: a scalable framework for spatial omics analysis. Nat. Methods 19, 171–178 (2022).

9. Wolf, F. A., Angerer, P. & Theis, F. J. SCANPY: large-scale single-cell gene expression data analysis. Genome Biol. 19, 15 (2018).

10. The MONAI Consortium. Project MONAI. (2020). doi:10.5281/zenodo.4323059.

11. Gayoso, A. et al. scvi-tools: a library for deep probabilistic analysis of single-cell omics data. Preprint at https://doi.org/10.1101/2021.04.28.441833.

12. Virshup, I. et al. The scverse project provides a computational ecosystem for single-cell omics data analysis. Nat. Biotechnol. (2023) doi:10.1038/s41587-023-01733-8.

13. Janesick, A. et al. High resolution mapping of the breast cancer tumor microenvironment using integrated single cell, spatial and in situ analysis of FFPE tissue. bioRxiv 2022.10.06.510405 (2022) doi:10.1101/2022.10.06.510405.

14. Gao, R. et al. Delineating copy number and clonal substructure in human tumors from single-cell transcriptomes. Nat. Biotechnol. 39, 599–608 (2021).

15. Wu, S. Z. et al. A single-cell and spatially resolved atlas of human breast cancers. Nat. Genet. 53, 1334–1347 (2021).

16. Kleshchevnikov, V. et al. Cell2location maps fine-grained cell types in spatial transcriptomics. Nat. Biotechnol. 40, 661–671 (2022).

17. Keller, M. S. et al. Vitessce: a framework for integrative visualization of multi-modal and spatially-resolved single-cell data. Preprint at https://doi.org/10.31219/osf.io/y8thv (2021).

18. ngff: Next-generation file format (NGFF) specifications for storing bioimaging data in the cloud. (Github).

19. Lohoff, T. et al. Integration of spatial and single-cell transcriptomic data elucidates mouse organogenesis. Nat. Biotechnol. (2021) doi:10.1038/s41587-021-01006-2.

20. van den Brink, S. C., et al. Single-cell and spatial transcriptomics reveal somitogenesis in gastruloids. Nature 582, 405–409 (2020).

21. Jackson, H. W. et al. The single-cell pathology landscape of breast cancer. Nature 578, 615–620 (2020).

22. Lin, J.-R. et al. Multiplexed 3D atlas of state transitions and immune interaction in colorectal cancer. Cell 186, 363–381.e19 (2023).

23. Irmisch, A. et al. The Tumor Profiler Study: integrated, multi-omic, functional tumor profiling for clinical decision support. Cancer Cell 39, 288–293 (2021).

24. Moses, L. & Pachter, L. Museum of spatial transcriptomics. Nat. Methods 19, 534–546 (2022).

25. Salmén, F. et al. Barcoded solid-phase RNA capture for Spatial Transcriptomics profiling in mammalian tissue sections. Nat. Protoc. 13, 2501–2534 (2018).

26. Eng, C.-H. L. et al. Transcriptome-scale super-resolved imaging in tissues by RNA seqFISH. Nature 568, 235–239 (2019).

27. Chen, K. H., Boettiger, A. N., Moffitt, J. R., Wang, S. & Zhuang, X. RNA imaging. Spatially resolved, highly multiplexed RNA profiling in single cells. Science 348, aaa6090 (2015).

28. He, S. et al. High-plex imaging of RNA and proteins at subcellular resolution in fixed tissue by spatial molecular imaging. Nat. Biotechnol. 40, 1794–1806 (2022).

29. McCormick, M. spatial-image/spatial-image: spatial-image 0.2.1. (2022). doi:10.5281/zenodo.6508869.

30. Hoyer, S. & Hamman, J. J. xarray: N-D labeled Arrays and Datasets in Python. J. Open Res. Softw. 5, (2017).

31. datatree: WIP implementation of a tree-like hierarchical data structure for xarray. (Github).

32. Gillies, S. et al. Shapely. (Zenodo, 2023). doi:10.5281/ZENODO.5597138.

33. Jordahl, K. et al. geopandas/geopandas: v0.8.1. (2020). doi:10.5281/zenodo.3946761.

34. Dask Development Team. Dask: Library for dynamic task scheduling. Preprint at https://dask.org (2016).

35. Virshup, I., Rybakov, S., Theis, F. J., Angerer, P. & Alexander Wolf, F. anndata: Annotated data. bioRxiv 2021.12.16.473007 (2021) doi:10.1101/2021.12.16.473007.

36. He, S., et al. High-Plex Multiomic Analysis in FFPE Tissue at Single-Cellular and Subcellular Resolution by Spatial Molecular Imaging. bioRxiv 2021.11.03.467020 (2021) doi:10.1101/2021.11.03.467020.

37. Schapiro, D. et al. MCMICRO: a scalable, modular image-processing pipeline for multiplexed tissue imaging. Nat. Methods 19, 311–315 (2022).

38. Long, B., Miller, J. & Consortium, T. S. T. SpaceTx: A Roadmap for Benchmarking Spatial Transcriptomics Exploration of the Brain. arXiv preprint arXiv:2301.08436 (2023).

39. Hartmann, F. J., et al. Multiplexed Single-cell Metabolic Profiles Organize the Spectrum of Cytotoxic Human T Cells. bioRxiv 2020.01.17.909796 (2020) doi:10.1101/2020.01.17.909796.

40. Eling, N. & Windhager, J. Example imaging mass cytometry raw data. (2022). doi:10.5281/zenodo.5949116.

41. Eling, N. & Windhager, J. steinbock results of IMC example data. (2022). doi:10.5281/zenodo.7412972.

42. Paszke, A., et al. Automatic Differentiation in PyTorch. in NIPS Autodiff Workshop (2017).

43. Erickson, A. et al. Spatially resolved clonal copy number alterations in benign and malignant tissue. Nature 608, 360–367 (2022).

